# Robust hierarchically organized whole-brain patterns of dysconnectivity in schizophrenia spectrum disorders observed after Personalized Intrinsic Network Topography

**DOI:** 10.1101/2022.12.13.520333

**Authors:** Erin W Dickie, Saba Shahab, Colin Hawco, Dayton Miranda, Gabrielle Herman, Miklos Argyelan, Jie Lisa Ji, Jerrold Jeyachandra, Alan Anticevic, Anil K. Malhotra, Aristotle N Voineskos

**Author notes:** Corresponding Author: Erin W Dickie, Centre for Addiction and Mental Health, 250 College Street, Rm 101, Toronto, Ontario, Canada, M5T 1R8.

## Abstract

**Background:** Spatial patterns of brain functional connectivity can vary substantially at the individual level. Applying cortical surface-based approaches with individualized rather than group templates may accelerate the discovery of biological markers related to psychiatric disorders. We investigated cortico-subcortical networks from multi-cohort data in people with schizophrenia spectrum disorders (SSDs) and healthy controls using individualized connectivity profiles.

**Methods:** We utilized resting state and anatomical MRI data from n=406 participants (n = 203 SSD, n = 203 healthy controls) from four cohorts. For each participant, functional timeseries were extracted from 80 cortical regions of interest, representing 6 intrinsic networks using 1) a volume-based approach 2) a surface-based group atlas approach, and 3) Personalized Intrinsic Network Topography (PINT), a personalized surface-based approach (Dickie et al., 2018). Timeseries were also extracted from previously defined intrinsic network subregions of the striatum (Choi et al 2011), thalamus (Ji et al 2019), and cerebellum (Buckner et al 2011).

**Results:** Compared to a volume-based approach, the correlations between all cortical networks and the expected subregions of the striatum, cerebellum, and thalamus were increased using a surface-based approach (Cohen’s D volume vs surface 0.27-1.00, all p<10^-6) and further increased after PINT (Cohen’s D surface vs PINT 0.18-0.96, all p <10^-4). In SSD vs HC comparisons, controlling for age, sex, scanner and in-scanner motion, we observed robust patterns of dysconnectivity that were strengthened using a surface-based approach and PINT (Number of differing pairwise-correlations: volume: 357, surface: 562, PINT: 630, FDR corrected). These patterns were found from four different cortical networks – frontal-parietal, sensory-motor, visual, and default mode -- to subcortical regions.

**Conclusion:** Our results indicate that individualized approaches can more sensitively delineate cortical network dysconnectivity differences in people with SSDs. These robust patterns of dysconnectivity were visibly organized in accordance with the cortical hierarchy, as predicted by computational models (Murray et al 2019). Our results also change our understanding of the specific network-network functional connectivity alterations in people with SSDs, and the extent of those alterations. Future work will examine these new patterns of dysconnectivity with behaviour using dimensional models.

**Highlights:**

- We evaluated the impact of cortical mapping method (volume-based, surface-based, vs surface personalized: PINT) on resting-state fMRI results in Schizophrenia Spectrum Disorders (SSD).
- The use of surface-based approaches and PINT increased the connectivity of cortical networks with the expected subregions of the striatum, thalamus and cerebellum, in comparison to a volume-based approach
- whole-brain case-control differences in functional connectivity were more pronounced after surface-based approach and PINT, in comparison to a volume-based approach

## Introduction

The search for the neurobiological underpinnings of schizophrenia spectrum disorders (SSDs) has been challenging, with no single brain system seemingly accounting for the diverse phenotypic expression of symptoms in people with an SSD. However, a few consistent findings have begun to emerge, including the notion of schizophrenia as a disorder of ‘dysconnectivity’ within the brain (Friston and Frith, 1995; Pettersson-Yeo et al., 2011). While findings in cortical-cortical connectivity have been mixed (Dong et al., 2018; Driesen et al., 2013; Li et al., 2019; Pettersson-Yeo et al., 2011; Whitfield-Gabrieli et al., 2009), connectivity between cortical and subcortical regions has emerged as a putative biomarker of SSD (Giraldo-Chica and Woodward, 2017; Ji et al., 2019b; Ramsay and MacDonald, 2018). Consistent patterns of dysconnectivity have been identified in the SSDs between cortical areas and the thalamus (Anticevic et al., 2015; Cao et al., 2018; Jacobs et al., 2019; Tu et al., 2018; Viviano et al., 2018; H.-L. S. Wang et al., 2015; Woodward et al., 2012), striatum (Avram et al., 2018; Ji et al., 2019b; Karcher et al., 2019; Li et al., 2020; Sarpal et al., 2015), and cerebellum (Brady et al., 2019; Dong et al., 2020; Ji et al., 2019a; Wang et al., 2014). The cortical networks implicated in these patterns are robust. Evidence of hyperconnectivity is observed for subcortical connectivity with visual and sensory motor regions and evidence of hypo-connectivity is observed for subcortical connectivity with fronto-parietal regions. These patterns of cortical-subcortical dysconnectivity, which reflect cortical hierarchy, are predicted with biophysical models of excitation/inhibition imbalance in psychosis (Murray et al., 2018).

Neuroimaging represents the primary approach to explore connectivity and brain function in SSD in vivo. Exploring these patterns of connectivity allows for the characterization of function within specific brain regions and across networks. However, the majority of research in psychiatry relies upon group averaging approaches using common templates derived from healthy control samples. This poses methodological challenges as considerable variability in the patterns of connectivity are present even amongst healthy individuals (Bijsterbosch et al., 2018; Gordon et al., 2017; D. Wang et al., 2015). This heterogeneity may be even greater in psychiatric populations, at both the structural and functional levels (Das et al., 2018; Dickie et al., 2018; Wang et al., 2018). Recent work has highlighted the possibility of greater “false positives” in biomarker research due to high levels of variability in clinical samples (Dickie et al., 2018; Hahamy et al., 2015). Better accounting for heterogeneity in clinical samples is therefore crucial to advance biomarker discovery and advance new treatment innovation.

Two important sources of individual variability are brain structure and functional topography. New approaches in neuroimaging are emerging to better account for these sources of variability. For example, as the human cortical surface is folded differently in all individuals, the use of 2-dimensional cortical surface-based anatomical registration may outperform 3-dimensional registration across individuals (Coalson et al., 2018; Glasser et al., 2016). These approaches can better account for variability in brain structure across individuals, but are not widely implemented across psychiatric neuroimaging. Even accounting for topographic variability in brain structure, there remains significant individual variability in functional topography; that is, regions of the brain which are related to specific cognitive processes or involved in different brain networks. Approaches which account for some variation on individualized functional topography can improve our ability to detect brain-behaviour relationships in both healthy participants (Bijsterbosch et al., 2018; Kong et al., 2018), and to predict symptoms in participants with SSD (Nawaz et al., 2021; Wang et al., 2018). For example, variation in the functional topography in the dorsolateral prefrontal cortex has been associated with the severity of negative symptoms in participants with SSD (Nawaz et al., 2021), and variability in the spatial pattern of task-evoked activity is associated with working memory and aging in SSD (Gallucci et al., 2022). Most work to date has focused on cortical-cortical resting-state connectivity, in many cases omitting data from key subcortical structures entirely.

The present study aimed to examine patterns of dysconnectivity in SSD, particularly between cortical and subcortical regions, while better accounting for individual variability in brain structural and functional topography. We investigated resting-state functional connectivity patterns in 203 participants with SSD and 203 pseudo-matched healthy control participants across four cohorts. For these datasets, we employed three methods for cortical brain mapping: a volume-based approach (the ‘standard’ analytical pipeline), a surface-based approach (better accounting for structural variability) as well as Personalized Intrinsic Network Topography (PINT), an individualized cortical brain mapping technique which accounts for some variation in functional topography. We then considered: 1) What is the impact of cortical brain mapping approach on measures of those cortical-subcortical circuits shown to be impacted by SSD? 2) What, in turn, is the impact of the cortical mapping approach on measures of dysconnectivity in these circuits?

## Methods

### Participants

Participants were recruited from several sites as part of various studies. In total, participants originated from four sites: 1) Centre for Addiction and Mental Health 2) Zucker Hillside Hospital (ZHH) 3) The Center for Biomedical Research Excellence (COBRE) (çetin et al., 2014), and 4) UCLA Consortium for Neuropsychiatric Phenomics LA5c Study (CNP)(Poldrack et al., 2016)(Poldrack et al 2016). Resting-state and anatomical MRI data were collected in all scenarios (see Supplemental Table 1 for MR parameters for each cohort). All sites had approval from their local research ethics boards, and all participants in all samples signed informed consent. Demographic details for each cohort are summarized in Table 1. Additional details for each cohort are as follows:

**Table 1:** Characteristics of patients and controls at time of scanning. Mean fd: mean framewise displacement of participant during functional scans, calculated with Nipype. Education data were not available for all participants and were calculated with the N available. (CAMH-SSD: 66 of 67 scores; COBRE-CTRL: 21 of 27 participants; COBRE-SSD: 16 of 27; ZHH-CTRL 85 of 104, ZZH-SSD 56 of 83)

| Cohort | CAMH |  | COBRE |  | CNP |  | ZHH |  |
| --- | --- | --- | --- | --- | --- | --- | --- | --- |
| Diagnosis | CTRL | SSD | CTRL | SSD | CTRL | SSD | CTRL | SSD |
| n | 41 | 67 | 27 | 22 | 31 | 31 | 104 | 83 |
| Sex = M (%) | 22 (53.7) | 40 (59.7) | 23 (85.2) | 19 (86.4) | 26 (83.9) | 24 (77.4) | 47(45.2) | 63 (75.9) |
| Age (years, mean (SD)) | 26.37 (6.67) | 32.19 (8.47) | 33.93 (9.66) | 29.55 (12.14) | 33.77 (8.94) | 35.23 (9.32) | 25.42 (6.33) | 25.86 (9.01) |
| Education (years, mean (SD)) | 15.95 (1.95) | 13.681 (2.04) | 14.602(1.8 5) | 13.273(1.4 4) | 15.35 (1.52) | 12.68 (1.49) | 14.904 (1.95) | 13.365 (2.79) |
| MRI mean FD (mm, mean (SD)) | 0.10 (0.04) | 0.12 (0.06) | 0.18 (0.04) | 0.21 (0.08) | 0.15 (0.08) | 0.18 (0.07) | 0.12 (0.04) | 0.13 (0.06) |

### CAMH cohort

We included participants between the ages 18 - 50. The Structured Clinical Interview for Axis I DSM-5 Disorders was used to diagnose individuals with schizophrenia spectrum disorders. Patients who could not provide voluntary consent or had a change in antipsychotic medications or decrement in functioning over the past 30 days were excluded. Control subjects were screened for DSM-5 disorders and were excluded if the subjects had a DSM Axis I disorder or their first-degree relatives were diagnosed with a schizophrenia spectrum disorder, other psychotic disorder, bipolar disorder, or depressive disorder with psychotic features. Exclusion criteria for all participants included a diagnosis of intellectual disability (as indexed by a WTAR score < 71), a history of a DSM-5 substance use disorder over the past 6 months, prior psychosurgery, Type 1 diabetes mellitus, unstable or debilitating medical illnesses, pregnancy, head injury, a neurological disease with extrapyramidal signs, or contraindications to an MRI scan. Non-English speakers were also excluded.

### ZHH cohort

Patients and controls (ages 15-65) were assessed with the Structured Clinical Interview for DSM-IV 2 Axis I Disorders (SCID). Patient diagnoses were determined via consensus among 3 senior clinicians and confidence in the accuracy of the agreed-upon diagnosis was assigned on a scale of 1-4 (4 denotes the highest confidence). Only patients with diagnoses assigned a high confidence rating (3 or 4) were included in the current study. In addition, only those healthy control subjects who had no history of an Axis I disorder were included in the current study. MRI exams were conducted at North Shore University Hospital on a single 3T scanner (GE Signa HDx; General Electric, Milwaukee, Wisconsin). All scans were reviewed by a radiologist and a member of the research team. Any scan with significant artifacts was repeated. A subset of participants were also included in longitudinal investigations, in these cases, the scan closest to available neurocognitive (IQ) data was selected for including in this study.

### COBRE cohort

Patients and controls were between the ages of 18 and 65. All participants were assessed with the Structured Clinical Interview for DSM-IV 2 Axis I Disorders (SCID). Control subjects were screened for DSM-5 disorders and were excluded if the subjects had a DSM-IV-TR Axis I disorder or any of their first-degree relatives had a psychotic disorder.

Exclusion criteria for all participants included a history of neurological disorder, mental retardation, severe head trauma with loss of consciousness, or substance abuse/dependence within the last year, or contraindications to an MRI scan. MRI scans were collected at 3-Tesla Siemens Trio scanner. Data were downloaded via SchizConnect database. http://schizconnect.org/.(Aine et al., 2017)

### CNP cohort

Participants were between the ages of 21-50, and had a primary language of English or Spanish. Patients and controls were assessed with the Structured Clinical Interview for DSM-IV 2 Axis I Disorders (SCID). All raters were assessed for a minimum 0.85 kappa for diagnostic accuracy. Patients with schizophrenia were excluded if they had comorbid bipolar disorder or ADHD. Exclusion criteria for all participants included a positive urinalysis for drugs of abuse, fewer than 8 years of formal education, left-handedness, pregnancy, or contraindications to an MRI scan. (Poldrack et al., 2016) https://www.nature.com/articles/sdata2016110

### MR Quality Assurance and participant matching

See Supplemental Table 2 for the breakdown of participant exclusions by cohort. Across the four cohorts, we started with 767 resting-state scans available from n=694 unique participants (n=317 SSD, n=377 HC). Scans from three participants were excluded for failing to have full brain coverage. Mean participant motion (framewise displacement (Power et al., 2014)) for resting state scans was calculated for each functional run using mriqc (Esteban et al., 2017). Resting-state scans were excluded if they had a mean framewise displacement greater than 0.5 or greater than 50% of frames with motion greater than 0.2mm per frame. If the remaining participants had multiple scans, one scan was selected per participant. For the ZHH data, if longitudinal data was available, the scan nearest the neurocognitive testing visit was chosen. For all other participants, the scan with the least motion was selected. Finally, due to a strong imbalance of diagnoses by cohort, we use the Machtit library in R to pseudo match participants so there were no group differences in age, sex and within scanner. For the CNP cohort - where a known ghosting artifact was recorded in a subsample of the resting-state scans - cases and controls were matched on age sex, scanner and presence of ghosting artifact. A final pseudo matched sample of 406 participants (n=203 SDD, n= 203) HC was carried forward for all further analyses.

### MR preprocessing

All T1w and fMRI images were preprocessed using a combination of the fMRIPREP ((Esteban et al., 2019a) and ciftify (Dickie et al., 2019) pipelines. FMRIprep is a Nipype (Gorgolewski et al., 2011, 2017) (RRID:SCR_002502) based tool. Each T1w (T1-weighted) volume was corrected for intensity non-uniformity using N4BiasFieldCorrection v2.1.0 (Tustison et al., 2010) and skull-stripped using antsBrainExtraction.sh v2.1.0 (using the OASIS template). Brain surfaces were reconstructed and subcortical regions were defined using recon-all from FreeSurfer v6.0.1(Dale et al., 1999) [RRID:SCR_001847]. Functional data was slice time corrected using 3dTshift from AFNI v16.2.07 (Cox, 1996) [RRID:SCR_005927] and motion corrected using mcflirt (FSL v5.0.9 (Jenkinson et al., 2002)). “Fieldmap-less” distortion correction was performed by co-registering the functional image to the same-subject T1w image with intensity inverted (Huntenburg, 2014; Wang et al., 2017) constrained with an average fieldmap template (Treiber et al., 2016), implemented with antsRegistration (ANTs). This was followed by co-registration to the corresponding T1w image using boundary-based registration(Greve and Fischl, 2009) with 9 degrees of freedom, using bbregister (FreeSurfer v6.0.1). Motion correcting transformations, field distortion correcting warp, BOLD-to-T1w transformation and T1w-to-template (MNI) warp were concatenated and applied in a single step using antsApplyTransforms (ANTs v2.1.0) using Lanczos interpolation. Many internal operations of FMRIPREP use Nilearn(Abraham et al., 2014) [ RRID:SCR_001362], principally within the BOLD-processing workflow. Following fMRIPREP, the ciftify workflows (Dickie et al., 2019) version 1.1.2-2.1.0 were used to convert the freesurfer reconstructed surfaces to gifti and cifti file formats. The cortical surfaces were realigned to the HCP fsLR templates (Glasser et al., 2013), using sulcal depth using the MSM algorithm (MSMSulc)(Robinson et al., 2014) and resampled to 32k vertices per hemisphere, and the freesurfer subcortical segmentation was used to define the participants 32k subcortical atlas greyordinates. The functional data was projected to the 32k surface coordinates using a ribbon constrained method that excludes outlier voxels, with methods similar to those employed by the HCP Pipelines (Glasser et al., 2013).

Three “dummy” TRs were removed from the beginning of each scan, and then the nilearn signal clean tool (Abraham et al., 2014) was used to apply a band-pass filter of 0.01-0.1 Hz, linear detrend, scale, and perform confound regression using six head motion parameters, white matter signal, CSF signal and global signal plus their lags, their squares, and the squares of their lags (i.e. a 24HMP + 8 Phys + 4 GSR confounds). Global signal regression was employed due to the benefits reported by (Ciric et al., 2018; Parkes et al., 2018). Confound regression was applied to both surface and volume data. To prevent the smoothing of nearby white matter signals into subcortical structures, no spatial smoothing was applied before timeseries extraction from the striatum, thalamus or cerebellum. Gaussian spatial smoothing with an 8mm FWHM was applied in the volume, before volume-based cortical timeseries extraction and on the cortical surface before template-based or personalized cortical surface data extraction (see below).

### fMRI Time Series Extraction and Connectivity Analysis

MNI coordinates for a series of 80 cortical ROIs were identified (Dickie et al., 2018), representing central nodes within subregions of six major resting-state networks (default mode, DM; fronto-parietal FP; ventral attention, VA; dorsal DA; sensory-motor, SM; visual, VI) based on the Yeo 7-network parcellation (Yeo et al., 2011). The cerebellum and striatum were parcellated according to parcellations by Buckner et al (2011) and Choi et al (2012). The thalamus was parcellated according to recent work by Ji et al. (2019a). In order to examine the impact of different preprocessing approaches in the examination of cortical to subcortical connectivity, functional connectivity matrices were derived using three analytical approaches (see Figure 1A below for schematic representation).

**Figure 1.**
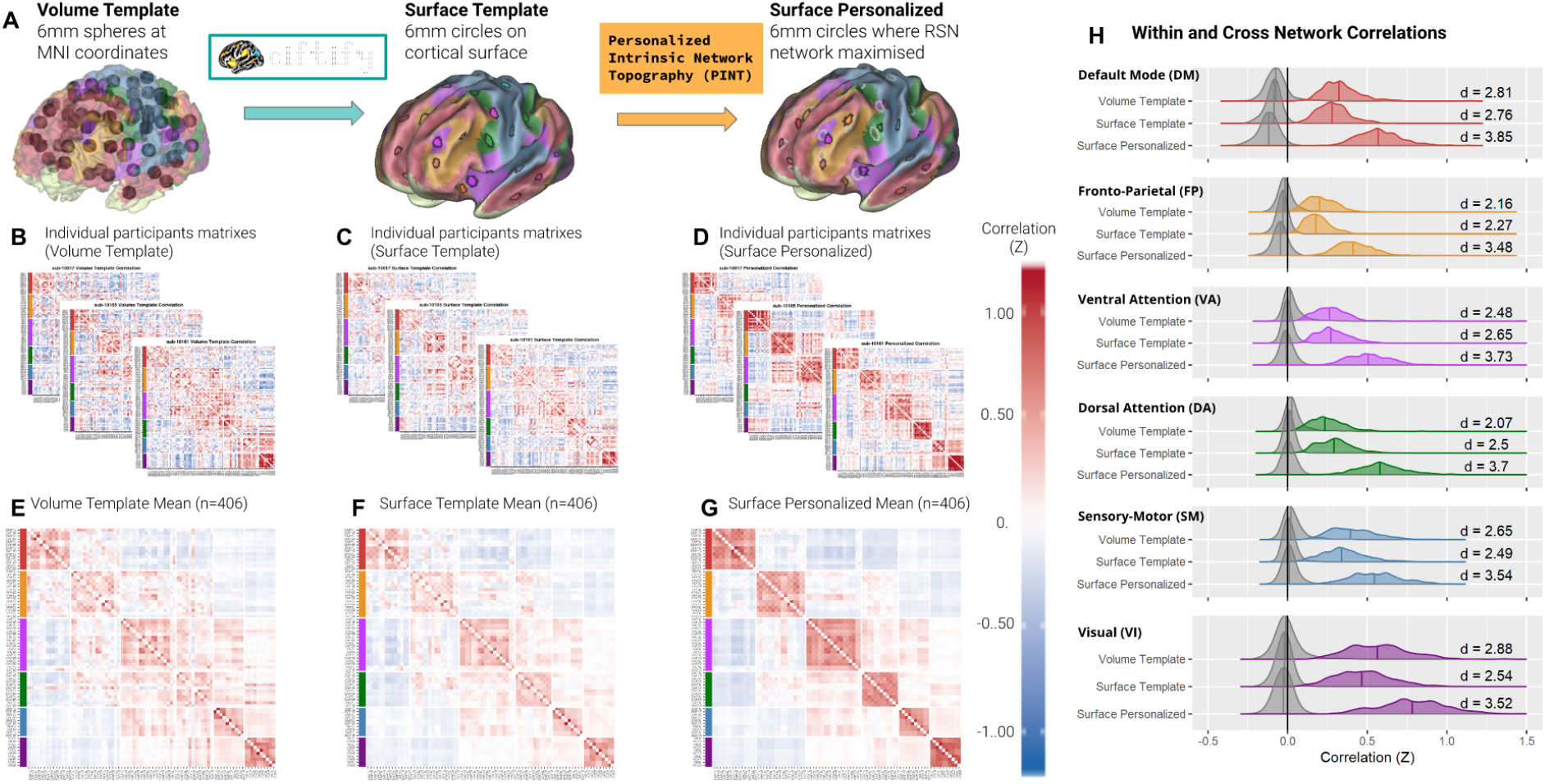
(A) Depiction of cortical mapping methods employed in this paper. In each case 80 cortical regions of interest were defined corresponding to 6 resting state networks of interest. Volume Template: Following anatomy based, 3D alignment to the MNI template, 6mm spheres surrounding MNI coordinates. Surface Template: Following anatomical 2D registration to the Conte surface template, using the ciftify workflow, 6mm circles are defined on the 2-dimensional cortical surface surrounding the same 80 points of interest. Surface Personalized: Using the PINT algorithm (Dickie et al., 2018), the 80 regions of interest are moved on the cortical surface to the location that maximize within-network resting-state connectivity in each network. (B-D) Cortical-cortical resting-state connectivity matrices from individual representative participants. (E-G) Mean (n=406) cortical-cortical resting state connectivity matrices. H. Distribution of within (in network color) and cross network (grey) resting state connectivity across participants using the three methods. Abbreviations: Abbreviations DM = Default mode, FP = Fronto-parietal, SM = Sensory Motor, VA = Ventral Attention.

**Method 1 - Volume-based analysis**: For each of the 80 cortical ROIs, a 6mm sphere was generated on the preprocessed volume-based fMRI data (i.e. prior to projection to the cortical surface).

**Method 2 - Surface-based Extraction:** For each of the 80 cortical ROIs, a 6mm circular ROI was generated on the cortical surface mesh.

**Method 3 - Personalized Intrinsic Network Topography (PINT):** Personalized cortical locations were defined for each participant using PINT (Dickie et al., 2018). PINT starts with the 80 surface-based 6mm cortical ROIs used in Method 2 above. Mean time series are extracted from each ROI, and the position of each ROI is then iteratively adjusted to maximize within-network connectivity. For each vertex within a given circular cortical ROI, the correlation of the vertex time series is calculated after the time series of the five other networks are regressed from both. Then, the center of the ROI is moved to the vertex of maximal partial correlation. After all 80 ROIs have been moved, the network time series are updated and the algorithm repeats around the new vertex locations. The algorithm iterates until all 80 ROIs are centered about a vertex of highest partial correlation. Average time series is then extracted from these 80 circular PINT ROIs (personalized ROIs accounting for individual functional network topography).

For each of the three methods, time series from each participant’s ROIs were correlated (Pearson’s R), then the correlation values were Z-transformed, to calculated edges of functional connectivity (FC). To characterize the effect of these cortical mapping algorithms, summary measures for within and between network connectivity were calculated for each participant for each of the 6 resting-state networks. Within network connectivity was calculated as the average Z-transformed correlation values for all ROIs within the same cortical network. Between-network connectivity was calculated for each network as the average of all edges from that cortical network to any of the five other cortical resting-state networks.

### Statistical Analysis

For all three cortical mapping techniques, for each edge (ROI to ROI FC), effects of SSD diagnosis was calculated in our sample of n=203 SSD participant and n=203 HC, using a linear model, with additional covariates of age, sex, site, and motion during the scan (mean framewise displacement). Multiple comparisons were corrected for using False Discovery Rate (FDR). For descriptive purposes, for each edge, we also calculated the Cohen’s D effect size for SSD cases vs controls, after “residualizing” FC values for the effects of age, sex, site, and motion during the scan.

To investigate these patterns further, edges were grouped according to cortical resting-state network (for cortical ROIs) and/or subcortical structure (striatum, thalamus, or cerebellum). Functional connectivity summary scores for these groups of edges were calculated for each participant by taking the weighted-mean of all edges that survived FDR correction (Cohen’s D threshold = 0.2678; means were weighted by the SSD diagnosis effect size). To account for the effect of education, case-control comparisons were re-run with education added as an additional covariate.

Given prior associations of cortical-subcortical connectivity with symptom scores and cognitive performance, we used these summary scores to explore associations with those clinical features we could characterise from these cohorts. For clinical symptom severity, we tested SANS for (CMH, UCLA, ZHH cohorts) and PANSS total scores (CAMH and COBRE cohorts).

Individual variability in intrinsic network locations was calculated as the geodesic distance from the starting “template” vertex location to each participant’s “personalized” vertex location during the PINT algorithm, measured on the HCP S1200 Average Surface in mm. This distance was averaged across all 80 ROIs for each participant to build a “total brain distance”, and within each network to produce network-level scores. We tested for effects of diagnosis and age on average distance using linear regression. All linear models included covariates of sex, cohort(site), total cortical surface area and mean framewise displacement during the fMRI resting-state scan.

All code related to the following analysis is available at https://github.com/TIGRLab/SSD_PINT.

## Results

### The use of a surface-based approach and PINT increased within network cortical-cortical connectivity

The sample mean cortical-cortical correlation matrices for fMRI time series extracted using a volume-based, surface-based and personalized surface-based (PINT) approach are shown in Figure 1. Within network correlations are higher than between network correlations, i.e. evidence of intrinsic network structure was observed for all networks tested, for the volume-based approach (Cohen’s D 2.07-2.88), as well as for the surface-based approach (Cohen’s D 2.27-2.76) and the personalized surface approach (Cohen’s D 3.48-3.85; see Figure 1H). Using a surface-based approach, as opposed to a volume-based approach, has a mixed effect on within-to-between network differences according to intrinsic network (see Supple Table 2): these within to between network correlation difference increase in the DA and VA networks, but decrease for the DM, SM, and VI networks. Using PINT increases within network correlations further for all networks (Surface Template vs PINT cohen D 2.41-3.10, see Suppl Table 2), but this is expected given that the PINT algorithm was explicitly designed to optimize these correlations.

### The use of surface-based approaches and PINT increased the connectivity of cortical networks with the expected subregions of the striatum, thalamus and cerebellum

In Figure 2, we plot the average correlations of four cortical networks (DM, SM, VA, FP) with previous parcellations of the striatum (Choi et al., 2012), thalamus (Ji et al., 2019a) and cerebellum (Buckner et al., 2011) (Note DA and VI networks were not tested because they are not well represented in the striatum). While cortical-subcortical correlations are noticeably weaker than the cortical-cortical correlations plotted in Figure 1, for all cortical timeseries extraction methods, for all four networks, we observed that the correlation to the expected subregions of the striatum, thalamus and cerebellum are positive, and greater than the correlations of these cortical networks to other subregions. Moreover, we observed that this expected pattern is strengthened when using a surface-based cortical timeseries extraction approach, as compared to the volume-based approach (cohen D (min-max) = 0.22-0.99). The pattern is again strengthened when using the surface-personalized approach (PINT), compared to the template surface-based approach (cohen D (min-max) = 0.14-0.96). In total, moving from a volume-based cortical timeseries extract methods to PINT increased the correlation with the expected cortical subregions by an effect size of 0.03-0.99, across the four cortical networks and subregions tested (see suppl Table 3). As shown in suppl table 3, this was a results of both increases in positive correlations with the expected subregions, as well as decreases in correlation (or increased anti-correlation) between networks. Note that PINT optimized within network connectivity based only on cortical ROIs, leaving subcortical regions fixed.

**Figure 2.**
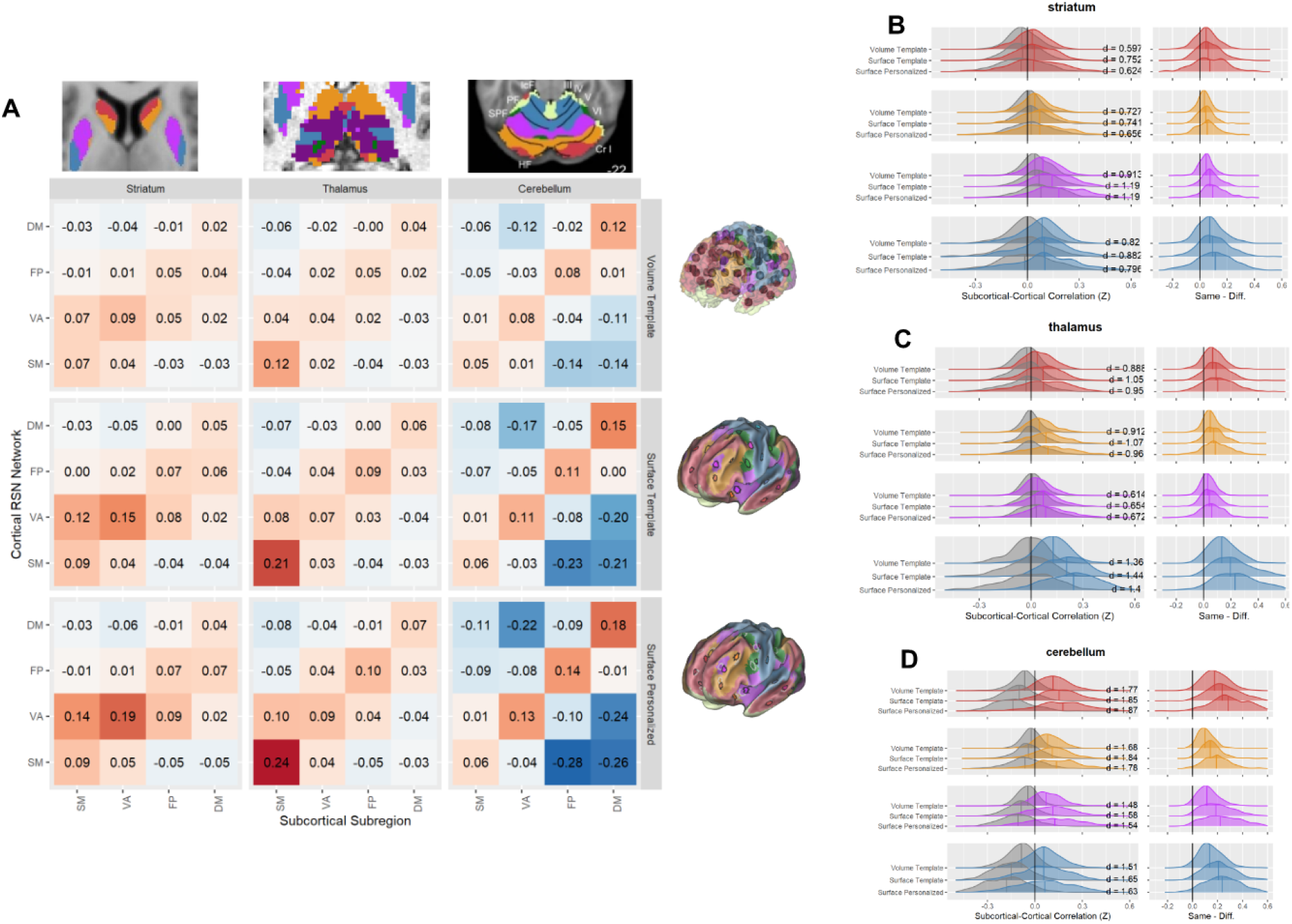
Cortical-subcortical functional connectivity as a function of cortical mapping method. A. Matrix of mean (n=406) functional correlation between cortical resting state networks and resting-state network associated subregions of the striatum, thalamus, and cerebellum. Cortical resting-state data was extracted using three methods (Volume Template, Surface Template and Surface Personalized - PINT (Dickie et al., 2018)). B-D On the left, the distribution (across) n=406 participants for same-network FC (left plot - in network color) and different-network FC (left-plot grey) and the same-different difference in FC (right-most plot, in network color). Where, B) Same network implies a match between the cortical resting-state network and resting state network associated subregions of the striatum defined in (Choi et al., 2012). c) Same network implies a match between the cortical resting-state network and resting state network associated subregions of the Thalamus defined in (Ji et al., 2019a). D) “Same network” implies a match between the cortical resting-state network and resting state network associated subregions of the Cerebellum define in (Buckner et al., 2011). Abbreviations DM = default mode, FP = Froto-parietal, SM = Sensory Motor, VA = Ventral Attention.

### Robust whole-brain profiles of dysconnectivity in SSD were more visible after surface-based approach and PINT

In figure 3, we plot edgewise effects of SSD diagnosis on functional connectivity, controlling for age, sex, site, and motion during the scan (mean framewise displacement). We observed robust patterns of dysconnectivity that were strengthened using a surface-based approach and PINT (Number of differing pairwise-correlations: volume: 419, surface-template: 579, surface-personalized (PINT): 644, FDR corrected). Moreover, patterns of dysconnectivity became more aligned with individual resting state networks and cortical hierarchy (see Figure 3D-E for breakdown by cortical network). Overall, regardless of cortical mapping approach, hypoconnectivity (decreased connectivity) was observed in participants with schizophrenia relative to controls for edges connecting subcortical regions to the FP network.

**Figure 3.**
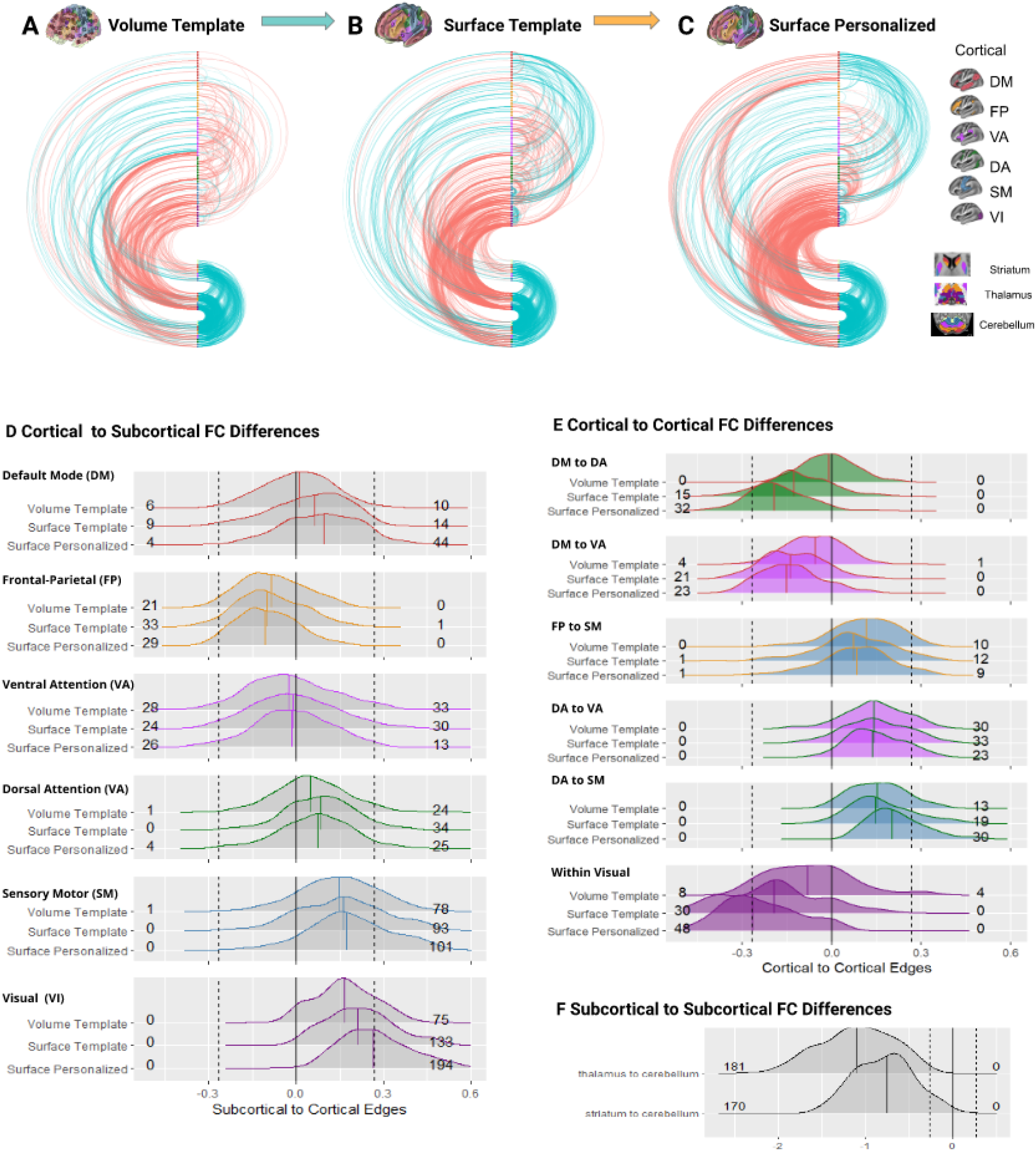
Edgewise SSD vs Healthy Control differences in resting-state functional connectivity. Resting-State connectivity edges showing significant (pFDR <0.05) hypoconnectivity (SSD < HC, in blue) and hyperconnectivity (HC > SSD, in red) are plotted as arcs, with opacity of arc is indicative of effect size. Brain regions of interest (ROIs) are marked by the circle ordered along the middle axis of each plot, with cortical regions of interest being in the top section - ordered by resting-state network followed by the Striatum, Thalamus and Cerebellum. Arcs are organized such that Cortical-cortical edges are plotted in the top right section, cortical-subcortical edges are plotted to the left, and subcortical-subcortical are plotted in the bottom right. Edges were significant in models that include Age, Sex, Scanner, framewise displacement as covariates. Notice that the number of significant edges increases as we move from A) Volume Template (n= 419 edges) to B) Surface Template (n= 579 edges) to C) Surface Personalized cortical mapping approach (n = 644 edges). In addition to more edges, the overall interpretability of the results in regards to resting state cortical networks also increase. D) The distributions of effect sizes of case-control differences are plotted for cortical-subcortical edges separately for each cortical-resting state network. The dotted lines on the plots mark the (pFDR < p.05 significance threshold) and the numbers to the left and right of each distribution mark the number of significant edges. E) The same types of distribution plots as (E) are created for the cortical-cortical resting-state networks most affected by cortical mapping method. F) Distribution of edgewise effect sizes for the subcortical-subcortical edges - note that these were kept constant as cortical mapping method varied.

Hyperconnectivity (increased connectivity) was observed in participants with SSD for subcortical connections to those VI, SM, and DA networks, which increased when moving from volume-to-surface, and again from surface-to-PINT. Additional patterns of hypoconnectivity where observed in the cortex between the DM network and the DA and VA networks, as well as within VI network edges. Cortical hyperconnectivity was observed between the DA and VA networks, as well as the FP network to the SM network. Hypoconnectivity was observed for nearly half of edges connecting the cerebellum to the striatum and the cerebellum to the thalamus.

To investigate these patterns further, sub-scores for these groups of edges were calculated for each participant by taking the weighted-mean of all edges that survived FDR correction (Cohen’s D threshold = 0.2678; means were weighted by the SSD diagnosis effect size). A significant main effect of SSD diagnosis was also observed when models were refitted with an additional covariate of years of Education (see Supplemental Table 4 for all model fits). When models were fit separately for each site all model fits remained significant for the two larger cohort ZHH and CAMH sample, with more variability observed in the COBRE and UCLA cohorts (see Supplemental Table 5 for all model fits).

Interestingly, when relationships were explored across different dysconnectivity patterns, hypoconnectivity between the subcortical areas and the FP network was negatively correlated with hyperconnectivity between subcortical areas and the VI network (see Supplemental Figure 4a).

### Relationships between connectivity and behavioral variables

Clinical scores (in SANS for CMH, UCLA, ZHH cohorts) and PANSS total scores (in CAMH and COBRE), were not correlated with any FC weighted scores after correction for multiple comparisons.

### Age and SSD diagnosis is associated with less variable network topography

We investigated the mean distance that ROIs travelled during the PINT algorithm as a measure of variability in intrinsic network topography. We used linear models with SSD diagnosis and age as predictors, and covariates of sex, site, total surface area and mean framewise displacement during the fMRI scan. For mean distance travelled, across all 80 ROIs, we observe a small effect of SSD diagnosis, such that participants with SSD had a smaller total distance travelled versus healthy controls (beta(SE)=-0.14(0.07), t=-2.05, p=0.041). Moreover, we observed a linear effect of age on ROI RSN network location variability, where older adults a smaller travel distance than younger adults (beta(SE)=0.57(0.16), t=3.51, p=0.0005, see Supplemental Figure 1b). When we tested mean distance travelled separately for each of 6 intrinsic networks, a significant effect of SSD diagnosis was not observed in any individual network after Bonferroni correction. The negative linear effect of age was observed for the visual (beta(SE)=1.14(0.41), t=2.77, p=0.035) and sensory motor networks (beta(SE)=1.63(0.54), t=3.00, p=0.017, see supplemental Figure 1c).

## Discussion

We examined the effects of utilizing a more individualized analytical framework on established cortical and subcortical dysconnectivity in SSD. We were able to create a large sample of participants by pooling data acquired across multiple studies/sites, matching within site on controls and SSD. We found that the most commonly used volumetric approach underestimated connectivity relative to the more ‘personalized’ approaches accounting for variability in anatomy and functional topography. The change was particularly pronounced from volume to surface analysis, which may in part be related to improvements in anatomical registration when using surface as opposed to volume based registration (Dickie et al., 2019). Moving from a volume-based to surface-based cortical mapping approach yielded stronger and more interpretable case-control differences in our samples of participants with SSD. These patterns were especially present in connectivity between cortical networks and the thalamus, striatum and cerebellum. Using a surface-based personalized approach (PINT) - as opposed to surface-based anatomical approach, yielded further refinements in the connectivity measures, especially cortical-subcortical connectivity, though the improvements were moderate compared to the volume to surface changes. However, the great cortical-subcortical dysconnectivity following PINT is noteworthy in that PINT works by optimizing connectivity within the cortical networks irrespective of connectivity with subcortical networks. These findings suggest that personalizing cortical ROIs impacts cortical-subcortical connectivity measures as well, potentially providing more accurate estimates on the individual level.

### PINT enhances and clarifies cortical-subcortical connectivity differences between people with an SSD and healthy controls

These results are in strong agreement with previous research that has shown robust patterns of dysconnectivity in SSD between cortical areas and the thalamus (Anticevic et al., 2015; Cao et al., 2018; Jacobs et al., 2019; Tu et al., 2018; Viviano et al., 2018; H.-L. S. Wang et al., 2015; Woodward et al., 2012), striatum (Avram et al., 2018; Ji et al., 2019b; Karcher et al., 2019; Li et al., 2020; Sarpal et al., 2015), and cerebellum (Brady et al., 2019; Dong et al., 2020; Ji et al., 2019a; Wang et al., 2014). Previous work has suggested these findings are heterogeneous - both hypo and hyper connectivity has been observed. However, when whole cortex patterns are examined, such as in the plots in Figure 3, it became clearer that patterns of subcortical cortical connectivity were organized according to cortical hierarchy, with evidence of hyperconnectivity observed for subcortical connectivity lower in cortical hierarchy (visual and sensory motor regions) and evidence of hypo-connectivity for subcortical connectivity with areas higher in cortical hierarchy (fronto-parietal regions). This pattern is predicted with biophysical models of excitation/inhibition imbalance in psychosis (Murray et al., 2018). Also, in keeping with these hypotheses, and replicating previous work(Avram et al., 2018; Ji et al., 2019b), we observed that dysconnectivity of subcortical areas to lower and higher regions of cortical hierarchy was correlated across participants, suggesting that both patterns may be markers of the same pathological process.

As many studies of improved cortical mapping methods have focused on cortical-cortical circuits, one area that has been less explored is the role that improving cortical mapping methods may have on measures of cortical-subcortical connectivity patterns. Our results suggest that improving cortical mapping methods, (i.e. using a surface-based approach, using personalized intrinsic mapping) can improve measures of these critical cortical-subcortical circuits. Previously described relationships between cortical resting state networks and specific subregions of the striatum, thalamus and cerebellum were more apparent after a surface based approach was used as opposed to a volume-based approach, and even stronger when a precision cortical mapping approach was employed. As the extracted subcortical data was kept constant across these three comparisons, we consider this as adding important evidence for the validity of these cortical mapping methods - in other words, evidence that the cortical networks these algorithms are behaving as expected. Furthermore, the strength of SSD case-control differences in cortical-subcortical networks was increased, as evidenced by more significant edges, as more advanced (i.e. surface-based, then personalized) mapping cortical techniques were applied. We suggest that these cortical methods have successfully reduced unwanted variance in functional connectivity due to individual heterogeneity in cortical anatomy and functional topography, leading to a more clinically relevant signal.

While methodologically robust patterns of dysconnectivity appear for cortical-subcortical connections in the brain, patterns of cortical-cortical connectivity were more sensitive to the cortical mapping method applied. It is possible that known heterogeneity in findings of cortical-cortical dysconnectivity across research groups may be, in-part, driven by differences in known mapping techniques(Dong et al., 2018; Fornito et al., 2012; Pettersson-Yeo et al., 2011). As we moved from a volume-based cortical approach to a surface-based approach, we saw an increase in the overall number of case-control differences and a stronger alignment of case-control differences with known resting-state cortical networks. In particular, by moving from volume-based to surface-based approaches, hypoconnectivity between the default-mode network and the dorsal attention network was observed. The use of a personalized mapping result seemed to lead to stronger congruence of results with resting state networks but not an increase in effect sizes. Therefore, while the benefits of moving from a traditional volume-based MR approach to the surface-based approach are clear, the benefits of using a personalized mapping method are less dramatic. We hope their value can be clarified as additional personalized mapping methods are introduced and evaluated.

### Looking at cortical network locations across individuals. Is there cortical “idiosyncrasy” in SSD?

The neurodevelopmental theory of schizophrenia (Owen et al., 2011) posits that SSD is a neurodevelopmental disorder that onsets in “later” brain development stages (i.e. late adolescence and early adulthood). Our previous work, using the same brain mapping approaches (Dickie et al., 2018), showed that ASD, another neurodevelopmental disorder associated with patterns of dysconnectivity, was associated with the “idiosyncratic” brain, i.e. increased variability in brain network topography(Dickie et al., 2018; Hahamy et al., 2015; Nunes et al., 2019). In this study, we did not see more variability in cortical network topography in participants with SSD compared to healthy controls. However, we should note that the PINT method is only one measure of idiosyncrasy in cortical development and other measures for differing cortical patterns in SSD have shown differences (Nawaz et al., 2021)(Das et al., 2018; Sun et al., 2020). Differences between our own results in SSD and our previous results in an ASD sample may relate to the differing developmental trajectories associated with their differing patterns of onset (Goldstein et al., 2002). Indeed, in both this study and our previous work (Dickie et al., 2018), we observed that individual differences in cortical network decrease with increasing age of the sample. The current study extended this work to include younger and older adults. Given these known effects of both clinical and developmental factors in cortical network organization, the use of surface-based and personalized mapping approaches becomes critical, as was seen in this paper, to understand clinical network properties. This is indeed supported by previous work showing benefits of surface-based (Anticevic et al., 2008) as well as personalized (Wang et al., 2018) approaches in SSD research.

### Are these patterns of functional connectivity specific to schizophrenia and do they change over time?

Using a large, pooled cross-sectional sample, we delineated robust patterns of dysconnectivity between participants with SSD and healthy controls. However, the establishment of the association of these patterns of dysconnectivity is only the beginning of many exciting unanswered research. Due to the retrospective pooling of our samples, our ability to incorporate additional clinical and cognitive measures was limited due to protocol differences across studies. Importantly, given that we only investigated SSD vs healthy control differences, the specificity of these findings to psychosis is not given. Indeed, altered cortical-subcortical connectivity is also seen in autism (Cerliani et al., 2015; Holiga et al., 2019), as well as other disorders (Tu et al., 2018) and has been related in some studies to a general P-factor (Elliott et al., 2018). Finally, an important avenue for research will be to understand how these patterns of functional connectivity may or may not change over time. When do these patterns develop? And are they altered by successful treatment of psychosis. Indeed, there is some evidence that some of these patterns may be present in participants before the onset of psychosis (Anticevic et al., 2015; Cao et al., 2018). There is also evidence that some patterns of brain connectivity change with successful treatment (Sarpal et al., 2015). In order to tease these mechanisms apart we will need to work more with large transdiagnostic longitudinal samples.

### Concluding remarks

This work suggests that our choice of neuroimaging analysis tools really do matter. Spatial patterns of brain functional connectivity can vary substantially at the individual level. Improving unwanted sources of variability via anatomical and functional alignment provides our models to more meaningfully patterns of variation in brain activity across people. The results of this paper suggest that surface-based approaches are not only possible in large clinical samples, but potentially necessary, to delineated results needed for clinical translation. It is therefore exciting that openly available tools for surface-based individualized connectomics are being developed and advanced every day (Bijsterbosch et al., 2018; Dickie et al., 2019; Esteban et al., 2019b; Glasser et al., 2013; Kong et al., 2018; D. Wang et al., 2015). If more data is available per participant, these approaches may be further strengthened through the incorporation of temporal “state” dynamics (Reinen et al., 2018). However, surface-based techniques are still more challenging to apply - as greater tooling and support is available to volume-based techniques and atlases. More education and simpler tooling will be needed for these techniques to be applied at a wider scale, for the benefit of psychiatry research.

## Acknowledgements

The CAMH dataset was collected with support from the Canadian Institutes of Health Research and NIH (ANV), CAMH Foundation, and NIMH (1/3 R01 MH102324). The COBRE imaging data and phenotypic information was collected and shared by the Mind Research Network and the University of New Mexico funded by a National Institute of Health Center of Biomedical Research Excellence (COBRE) grant 1P20RR021938-01A2. The CNP was one of 9 Interdisciplinary Research Consortia supported by the NIH Roadmap Initiative from 2007–2012. The CNP comprised 8 linked awards, including a Coordinating Center (UL1DE019580, UL1RR024911), five linked R01 awards (RL1MH083268, RL1MH083269, RL1DA024853, RL1MH083270, and RL1LM009833) and two center grant awards (PL1MH083271 and PL1NS062410). ZHH data set was collected via support from NIMH grants P50MH080173 and RO1MH102313 (PI: Malhotra).

EWD receives funding from the Brain and Behavior Research Foundation Young Investigator Award (NARSAD), the Canadian Institutes of Health Research. A.N.V. receives funding from the National Institute of Mental Health, Canadian Institutes of Health Research, Canada Foundation for Innovation, CAMH Foundation, and the University of Toronto. AA receives funding from NIH grants DP5OD012109 (AA), 1U01MH121766 (AA), 5R01MH112189 (AA), 5R01MH108590 (AA), NIAAA grant 2P50AA012870-11 (AA), the Brain and Behavior Research Foundation Young Investigator Award (AA), SFARI Pilot Award (AA).

## Data and Code Availability Statement

All code related to the preprocessing and statistical analysis is available at https://github.com/TIGRLab/SSD_PINT. The preprocessing makes use of publicly available pipelines, namely FMRIprep (version 1.1.2, https://fmriprep.org/en/stable/) and ciftify (version 2.1.0, https://edickie.github.io/ciftify/). This analysis used a combination of locally shared and publicly available data. The COBRE cohort was data were downloaded via SchizConnect database. http://schizconnect.org/.(Aine et al., 2017). The CNP cohort data is available on Openneuro (https://openneuro.org/datasets/ds000030). Original scans from CAMH and ZHH dataset participants can be obtained through collaborative agreement and reasonable request but are not publicly available due to the lack of informed consent by these human participants. However, anolymized FC derivatives from all participants included in the final analysis are linked to the https://github.com/TIGRLab/SSD_PINT repository.

