## Supplemental Material for "Robust hierarchically organized whole-brain patterns of dysconnectivity in schizophrenia spectrum disorders observed after Personalized Intrinsic Network Topography"

Supplemental Table 1 - MRI scan characteristics by Site.

| Measure | CAMH | ZHH | COBRE | CNP |
|---|---|---|---|---|
| Recruitment age | 18-50 | 15-65 | 18-65 | 21–50 |
| Scanner | GE 3T Discovery MR750<br>8-channel head coil | GE Signa HDx | Siemens Tim-Trio 3T scanner | 3T Siemens Trio |
| T1w Parameters | FSPGR-BRAVO<br><br>TE/TR = 3/6.7ms<br><br>0.9 x 0.9 x 0.9 $mm^3$ | IR-FSPGR<br>TR = 7.5 ms, TE = 3 ms, TI = 650 ms matrix = 256x256, FOV = 240 mm<br>216 contiguous images (slice thickness = 1mm) | 3D MPRAGE sequence (TR/TE/TI=2530/[1.64, 3.5, 5.36, 7.22, 9.08]/900, flip angle=7°, voxel size (isotropic)=1mm, image size=256×256×176 voxels) | MPRAGE<br>TR=1.9 s, TE=2.26 ms, FOV=250 mm, matrix=256×256, sagittal plane, slice thickness=1 mm, 176 slices. |
| fMRI parameters | EPI<br>TE=30.0 ms, FOV = 20 cm, and a 90o flip angle<br>40 slices parallel to the axial plane | EPI<br>TE = 30 ms, 40 continuous axial oblique slices | T2*-weighted gradient-echo, echo-planar sequence, 32 axial slices parallel to the AC-PC using a (TE=29ms, flip angle=75°, | T2*-weighted echoplanar imaging (EPI)slice,TE=30 ms, flip angle=90°, oblique slice orientation |
| BOLD matrix | 64x64<br>40 slices | 64x64<br>40 slices | 64x64<br>32 slices | 64x64<br>34 slices |
| BOLD resolution | 3.125 x 3.125 x 4 mm | 3.75x3.75x3 mm | 3×3×4mm | 3.75x3.75x4 mm |
| BOLD TR | 2 seconds | 2 seconds | 2 seconds | 2 seconds |
| BOLD length | 208 frames<br>6:56 | 148 frames<br>4:56 | 149 frames<br>4:58 | 152 frames<br>5:04 |
| Resting State | Eyes Closed | Eyes Closed | Eyes Open | Eyes Open |

Supplemental Table 2: Sample sizes after quality assurance steps.

| Site | CMH | | ZHH | | COBRE | | CNP ds000030 | |
|---|---|---|---|---|---|---|---|---|
| DX | CTRL | SSD | CTRL | SSD | CTRL | SSD | CTRL | SSD |
| original number of scans | 42 | 73 | 124 | 175 | 90 | 91 | 122 | 50 |
| original number of unique subjects | 41 | 67 | 124 | 115 | 90 | 85 | 122 | 50 |
| number subject excluded for not having full head coverage (i.e. cerebellum cut off) during fMRI scan | 0 | 0 | 2 | 1 | 0 | 0 | 0 | 0 |
| number of subjects excluded for motion in fMRI scan(s)* | 1 | 6 | 12 | 31 | 55 | 63 | 15 | 19 |
| number excluded in the process of matching | 0 | 0 | 6 | 0 | 8 | 0 | 76 | 0 |
| final number of participants | 41 | 67 | 104 | 83 | 27 | 22 | 31 | 31 |

*Exclusion for motion meaning no more than one scans that meant criteria. Criteria being a mean framewise displacement under 0.5 and less than 50% of frames with motion greater than 0.2mm per frame. (as established by MRIQC, 0.10.4 (Esteban et al., 2017)).

**Supplemental Figure 1**. Association of intrinsic network variability with SSD Diagnosis and Age. A) Distances travelled during the PINT algorithm (mm) averaged across all 80 regions of interest (ROIs) are plotted in black for healthy participants and red for participants with SSD across by cohort. B) Residuals PINT distance travelled (values from y-axis of figure A) were residualized for effects of Site (i.e. cohort), sex, total surface area, and motion during the fMRI scan (framewise displacement), and plotted vs Age (on the x-axis, (beta(SE)=0.57(0.16), t=3.51, p=5e-04) and diagnosis (healthy controls in black and participants with SSD in red, (beta(SE)=-0.14(0.07), t=-2.05, p=0.041)). C) The relationship between Age, Sex and PINT distance travelled is plotted separately for each intrinsic network, after controlling (residuals) for the same covariates. A significant effect of SSD diagnosis was not observed in any individual network after Bonferroni correction for the 6 networks tested. The negative linear effect of age was observed for the visual (beta(SE)=1.14(0.41), t=2.77, p=0.035) and sensory-motor networks (beta(SE)=1.63(0.54), t=3.00, p=0.017)

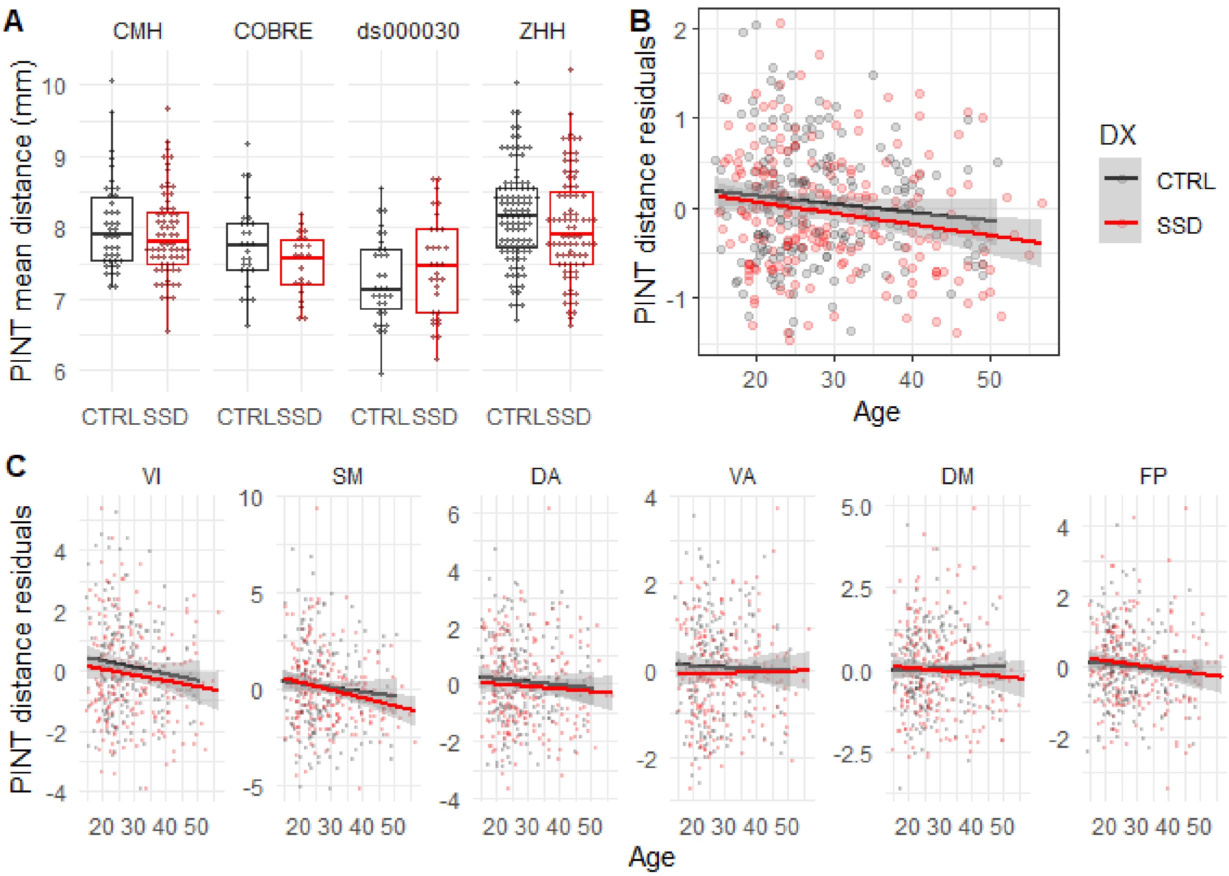

Supplemental Table 3. Changes (Cohen D) and paired t-test comparing for cortical-cortical correlations across different cortical surface data extraction methods (n = 406).

| Cortical Intrinsic Network | Change in Within Network Correlation | Change in Cross Network Correlation | Change in Within-Cross Network Correlation |
|---|---|---|---|
| ***Comparing Template Surface to Template Volume*** | | | |
| DM | d = -0.46, t(405) =-9.24, p= 8.6e-18 | d = -0.29, t(405) =-5.80, p=8.0e-08 | d = -0.20, t(405) =-3.96, p= 5.3e-04 |
| FP | d = -0.29, t(405) =-5.84, p= 6.5e-08 | d = -0.32, t(405) =-6.35, p=3.5e-09 | d = 0.01, t(405) =0.13, p=ns |
| VA | d = 0.09, t(405) =1.88, p=ns | d = -0.34, t(405) =-6.92, p=1.1e-10 | d = 0.57, t(405) =11.50, p= 6.9e-26 |
| DA | d = 0.32, t(405) =6.42, p= 2.3e-09 | d = -0.36, t(405) =-7.26, p=1.2e-11 | d = 0.86, t(405) =17.35, p= 5.0e-50 |
| SM | d = -0.52, t(405) =-10.42, p= 6.6e-22 | d = -0.31, t(405) =-6.35, p=3.5e-09 | d = -0.40, t(405) =-8.10, p= 3.8e-14 |
| VI | d = -0.75, t(405) =-15.04, p= 3.6e-40 | d = -0.28, t(405) =-5.54, p=3.2e-07 | d = -0.57, t(405) =-11.54, p= 4.8e-26 |
| ***Comparing Template Surface to Personalized Surface*** | | | |
| DM | d = 3.26, t(405) =65.75, p=1.9e-217 | d = -0.97, t(405) =-19.46, p=3.1e-59 | d = 3.10, t(405) =62.51, p=2.3e-209 |
| FP | d = 3.13, t(405) =63.14, p=6.1e-211 | d = -0.54, t(405) =-10.90, p=1.1e-23 | d = 2.84, t(405) =57.15, p=3.4e-195 |
| VA | d = 3.04, t(405) =61.22, p=4.9e-206 | d = -0.22, t(405) =-4.45, p=6.6e-05 | d = 2.89, t(405) =58.24, p=3.8e-198 |
| DA | d = 2.86, t(405) =57.63, p=1.7e-196 | d = -0.13, t(405) =-2.65, p=0.049 | d = 2.72, t(405) =54.80, p=1.2e-188 |
| SM | d = 2.56, t(405) =51.60, p=2.2e-179 | d = 0.11, t(405) =2.20, p=ns | d = 2.41, t(405) =48.61, p=2.4e-170 |
| VI | d = 2.73, t(405) =54.97, p=4.1e-189 | d = -0.08, t(405) =-1.58, p=ns | d = 2.70, t(405) =54.48, p=9.9e-188 |

P values are bonferonni corrected across 6 networks.

Supplemental Table 4: Changes (Cohen D) and paired t-test comparing for subcortical-cortical correlations across different cortical surface data extraction methods (n = 406).

| Cortical Intrinsic Network | Change in Within Network Correlation | Change in Cross Network Correlation | Change in Within-Cross Network Correlation |
|---|---|---|---|
| ***Comparing Template Surface to Template Volume*** | | | |
| ***Cortical networks to Striatum*** | | | |
| DM | d = 0.32, t(405) =6.37, p=5.2e-10 | d = 0.05, t(405) =1.06, p=3.465 | d = 0.31, t(405) =6.24, p=1.1e-09 |
| FP | d = 0.27, t(405) =5.47, p=7.9e-08 | d = 0.20, t(405) =4.13, p=4.5e-05 | d = 0.22, t(405) =4.40, p=1.4e-05 |
| VA | d = 0.75, t(405) =15.14, p=2.5e-41 | d = 0.36, t(405) =7.32, p=1.3e-12 | d = 0.74, t(405) =14.94, p=1.7e-40 |
| SM | d = 0.17, t(405) =3.50, p=0.006 | d = -0.09, t(405) =-1.78, p=0.910 | d = 0.35, t(405) =7.01, p=9.9e-12 |
| ***Cortical networks to Thalamus*** | | | |
| DM | d = 0.29, t(405) =5.76, p=1.7e-08 | d = -0.08, t(405) =-1.54, p=1.493 | d = 0.41, t(405) =8.19, p=3.5e-15 |
| FP | d = 0.47, t(405) =9.39, p=4.4e-19 | d = 0.14, t(405) =2.86, p=0.054 | d = 0.51, t(405) =10.21, p=6.4e-22 |
| VA | d = 0.41, t(405) =8.30, p=1.6e-15 | d = 0.22, t(405) =4.52, p=8.0e-06 | d = 0.29, t(405) =5.94, p=6.0e-09 |
| SM | d = 0.89, t(405) =17.93, p=2.3e-53 | d = 0.04, t(405) =0.72, p=5.636 | d = 0.82, t(405) =16.61, p=1.2e-47 |
| ***Cortical networks to Cerebellum*** | | | |
| DM | d = 0.46, t(405) =9.32, p=7.4e-19 | d = -0.51, t(405) =-10.23, p=5.2e-22 | d = 0.84, t(405) =16.92, p=5.7e-49 |
| FP | d = 0.38, t(405) =7.66, p=1.4e-13 | d = -0.28, t(405) =-5.72, p=2.0e-08 | d = 0.80, t(405) =16.15, p=1.2e-45 |
| VA | d = 0.43, t(405) =8.64, p=1.3e-16 | d = -0.69, t(405) =-13.86, p=5.0e-36 | d = 0.99, t(405) =20.03, p=1.6e-62 |
| SM | d = 0.04, t(405) =0.81, p=5.011 | d = -0.84, t(405) =-17.03, p=2e-49 | d = 0.78, t(405) =15.62, p=2.2e-43 |
| ***Comparing Template Surface to Personalized Surface*** | | | |
| ***Cortical networks to Striatum*** | | | |
| DM | d = -0.05, t(405) =-0.93, | d = -0.19, t(405) =-3.86, | d = 0.14, t(405) =2.72, |

| | | | |
|---|---|---|---|
| | p=4.261 | p=0.002 | p=0.081 |
| FP | d = 0.07, t(405) =1.32, p=2.262 | d = -0.07, t(405) =-1.51, p=1.595 | d = 0.22, t(405) =4.35, p=1.7e-05 |
| VA | d = 0.64, t(405) =12.81, p=9.3e-32 | d = 0.28, t(405) =5.66, p=2.8e-08 | d = 0.65, t(405) =13.04, p=1.0e-32 |
| SM | d = 0.05, t(405) =1.04, p=3.602 | d = -0.09, t(405) =-1.77, p=0.932 | d = 0.18, t(405) =3.58, p=0.005 |
| ***Cortical networks to Thalamus*** | | | |
| DM | d = 0.07, t(405) =1.37, p=2.075 | d = -0.22, t(405) =-4.49, p=9.4e-06 | d = 0.33, t(405) =6.74, p=5.3e-11 |
| FP | d = 0.21, t(405) =4.27, p=2.4e-05 | d = -0.01, t(405) =-0.16, p=10.512 | d = 0.29, t(405) =5.84, p=1.1e-08 |
| VA | d = 0.42, t(405) =8.46, p=4.9e-16 | d = 0.19, t(405) =3.87, p=0.001 | d = 0.38, t(405) =7.63, p=1.7e-13 |
| SM | d = 0.52, t(405) =10.53, p=4.5e-23 | d = -0.03, t(405) =-0.54, p=7.075 | d = 0.55, t(405) =11.06, p=5.0e-25 |
| ***Cortical networks to Cerebellum*** | | | |
| DM | d = 0.31, t(405) =6.19, p=1.5e-09 | d = -0.74, t(405) =-14.92, p=2.0e-40 | d = 0.96, t(405) =19.35, p=1.5e-59 |
| FP | d = 0.42, t(405) =8.44, p=5.9e-16 | d = -0.41, t(405) =-8.35, p=1.1e-15 | d = 0.87, t(405) =17.51, p=1.7e-51 |
| VA | d = 0.33, t(405) =6.67, p=8.2e-11 | d = -0.53, t(405) =-10.59, p=2.8e-23 | d = 0.80, t(405) =16.03, p=3.9e-45 |
| SM | d = 0.00, t(405) =-0.02, p=11.771 | d = -0.66, t(405) =-13.31, p=9e-34 | d = 0.60, t(405) =12.04, p=9.9e-29 |

Supplemental Table 4. Main effect of SSD diagnosis of functional connectivity after including years of education as an additional model covariate. Additional model covariates included age, sex, site, and motion in the scanner.

| | Volume Template | Surface Template | Surface Personalized |
|---|---|---|---|
| Subcortical to Cortical DM | beta(SE)=0.06(0.01), t(336)=5.23, p=4.2e-06 | beta(SE)=0.07(0.01), t(336)=5.31, p=2.8e-06 | beta(SE)=0.08(0.01), t(336)=5.39, p=1.9e-06 |
| Subcortical To Cortical FP | beta(SE)=-0.06(0.01), t(336)=-6.09, p=4.4e-08 | beta(SE)=-0.08(0.01), t(336)=-6.09, p=4.4e-08 | beta(SE)=-0.08(0.01), t(336)=-6.26, p=1.6e-08 |
| Subcortical to Cortical VA | beta(SE)=-0.06(0.01), t(336)=-6.73, p=1.0e-09 | beta(SE)=-0.07(0.01), t(336)=-6.15, p=3.1e-08 | beta(SE)=-0.08(0.01), t(336)=-5.45, p=1.4e-06 |
| Subcortical to Cortical DA | beta(SE)=0.05(0.01), t(336)=4.98, p=1.4e-05 | beta(SE)=0.07(0.01), t(336)=4.91, p=2.0e-05 | beta(SE)=0.07(0.02), t(336)=3.94, p=0.001 |
| Subcortical to Cortical SM | beta(SE)=0.07(0.01), t(336)=6.60, p=2.3e-09 | beta(SE)=0.09(0.01), t(336)=6.23, p=1.9e-08 | beta(SE)=0.09(0.02), t(336)=5.42, p=1.6e-06 |
| Subcortical to Cortical VI | beta(SE)=0.06(0.01), t(336)=5.58, p=7.1e-07 | beta(SE)=0.08(0.01), t(336)=6.07, p=4.9e-08 | beta(SE)=0.09(0.02), t(336)=5.63, p=5.4e-07 |
| Cortical DM to Cortical VA | beta(SE)=-0.13(0.02), t(336)=-6.16, p=2.9e-08 | beta(SE)=-0.10(0.01), t(336)=-7.04, p=1.5e-10 | beta(SE)=-0.11(0.02), t(336)=-5.67, p=4.4e-07 |
| Cortical DM to Cortical DA | no edges | beta(SE)=-0.08(0.01), t(336)=-5.84, p=1.7e-07 | beta(SE)=-0.08(0.02), t(336)=-5.03, p=1.1e-05 |
| Cortical FP to Cortical SM | beta(SE)=0.09(0.02), t(336)=4.01, p=0.001 | beta(SE)=0.08(0.01), t(336)=5.14, p=6.4e-06 | beta(SE)=0.08(0.02), t(336)=4.33, p=2.8e-04 |
| Cortical VA to Cortical DA | beta(SE)=0.09(0.02), t(336)=5.17, p=5.7e-06 | beta(SE)=0.09(0.01), t(336)=6.60, p=2.2e-09 | beta(SE)=0.09(0.02), t(336)=4.70, p=5.4e-05 |
| Cortical DA to Cortical SM | beta(SE)=0.09(0.02), t(336)=4.30, p=3.2e-04 | beta(SE)=0.09(0.02), t(336)=5.92, p=1.1e-07 | beta(SE)=0.10(0.02), t(336)=5.54, p=8.3e-07 |
| Cortical VI to Cortical VI | beta(SE)=-0.10(0.03), t(336)=-3.57, p=0.006 | beta(SE)=-0.12(0.02), t(336)=-5.24, p=4.0e-06 | beta(SE)=-0.13(0.03), t(336)=-5.01, p=1.2e-05 |
| Cerebellum to Striatum | beta(SE)=-0.08(0.02), t(336)=-4.78, p=3.7e-05 | | |
| Cerebellum to Thalamus | beta(SE)=-0.11(0.02), t(336)=-6.47, p=4.9e-09 | | |

Supplemental Table 5. Main effect of SSD diagnosis of functional connectivity in models fit separately per cohort.. Additional model covariates included age, sex, site, and motion in the scanner.

| | Cohort | Volume Template | Surface Template | Surface Personalized |
|---|---|---|---|---|
| Subcortical to Cortical DM | CAMH | beta(SE)=0.05(0.02), t(103)=3.17, p=0.028 | beta(SE)=0.07(0.02), t(103)=3.73, p=0.004 | beta(SE)=0.05(0.02), t(103)=2.44, p=ns |
| Subcortical to Cortical DM | ZHH | beta(SE)=0.04(0.01), t(182)=3.11, p=0.031 | beta(SE)=0.07(0.01), t(182)=4.92, p=2.7e-05 | beta(SE)=0.09(0.02), t(182)=5.09, p=1.3e-05 |
| Subcortical to Cortical DM | COBRE | beta(SE)=0.05(0.02), t(44)=2.39, p=ns | beta(SE)=0.05(0.03), t(44)=1.74, p=ns | beta(SE)=0.07(0.03), t(44)=2.40, p=ns |
| Subcortical to Cortical DM | UCLA | beta(SE)=0.09(0.02), t(57)=4.37, p=7.4e-04 | beta(SE)=0.08(0.03), t(57)=2.87, p=0.080 | beta(SE)=0.11(0.03), t(57)=4.05, p=0.002 |
| Subcortical To Cortical FP | CAMH | beta(SE)=-0.07(0.02), t(103)=-4.18, p=8.7e-04 | beta(SE)=-0.07(0.02), t(103)=-3.46, p=0.011 | beta(SE)=-0.07(0.02), t(103)=-3.50, p=0.010 |
| Subcortical To Cortical FP | ZHH | beta(SE)=-0.05(0.01), t(182)=-4.68, p=7.8e-05 | beta(SE)=-0.07(0.02), t(182)=-4.77, p=5.2e-05 | beta(SE)=-0.07(0.02), t(182)=-4.43, p=2.3e-04 |
| Subcortical To Cortical FP | COBRE | beta(SE)=-0.04(0.02), t(44)=-2.22, p=ns | beta(SE)=-0.07(0.03), t(44)=-2.67, p=ns | beta(SE)=-0.09(0.03), t(44)=-3.00, p=0.061 |
| Subcortical To Cortical FP | UCLA | beta(SE)=-0.04(0.02), t(57)=-2.26, p=ns | beta(SE)=-0.08(0.02), t(57)=-3.18, p=0.033 | beta(SE)=-0.04(0.02), t(57)=-1.84, p=ns |
| Subcortical to Cortical VA | CAMH | beta(SE)=-0.06(0.01), t(103)=-4.01, p=0.002 | beta(SE)=-0.06(0.02), t(103)=-3.33, p=0.017 | beta(SE)=-0.05(0.02), t(103)=-2.40, p=ns |
| Subcortical to Cortical VA | ZHH | beta(SE)=-0.06(0.01), t(182)=-6.08, p=9.5e-08 | beta(SE)=-0.08(0.02), t(182)=-5.35, p=3.6e-06 | beta(SE)=-0.09(0.02), t(182)=-4.83, p=4.0e-05 |
| Subcortical to Cortical VA | COBRE | beta(SE)=-0.04(0.02), t(44)=-2.02, p=ns | beta(SE)=-0.06(0.03), t(44)=-2.43, p=ns | beta(SE)=-0.10(0.03), t(44)=-2.97, p=0.067 |
| Subcortical to Cortical VA | UCLA | beta(SE)=-0.04(0.01), t(57)=-3.23, p=0.029 | beta(SE)=-0.04(0.02), t(57)=-1.84, p=ns | beta(SE)=-0.05(0.03), t(57)=-1.63, p=ns |
| Subcortical to Cortical DA | CAMH | beta(SE)=0.08(0.02), t(103)=4.65, p=1.4e-04 | beta(SE)=0.07(0.02), t(103)=3.56, p=0.008 | beta(SE)=0.08(0.02), t(103)=3.40, p=0.014 |
| Subcortical to Cortical DA | ZHH | beta(SE)=0.06(0.01), t(182)=4.33, p=3.4e-04 | beta(SE)=0.08(0.02), t(182)=4.71, p=6.8e-05 | beta(SE)=0.08(0.02), t(182)=3.44, p=0.010 |
| Subcortical to Cortical DA | COBRE | beta(SE)=0.08(0.02), t(44)=3.66, p=0.009 | beta(SE)=0.11(0.04), t(44)=2.91, p=0.080 | beta(SE)=0.07(0.04), t(44)=1.54, p=ns |
| Subcortical to Cortical DA | UCLA | beta(SE)=0.03(0.02), t(57)=1.61, p=ns | beta(SE)=0.04(0.02), t(57)=1.95, p=ns | beta(SE)=0.04(0.03), t(57)=1.23, p=ns |

| | | | | |
|---|---|---|---|---|
| Subcortical to Cortical SM | CAMH | beta(SE)=0.07(0.02), t(103)=4.06, p=0.001 | beta(SE)=0.09(0.02), t(103)=4.37, p=4.2e-04 | beta(SE)=0.09(0.02), t(103)=3.74, p=0.004 |
| Subcortical to Cortical SM | ZHH | beta(SE)=0.05(0.01), t(182)=4.32, p=3.6e-04 | beta(SE)=0.07(0.02), t(182)=4.09, p=9.0e-04 | beta(SE)=0.07(0.02), t(182)=3.65, p=0.005 |
| Subcortical to Cortical SM | COBRE | beta(SE)=0.08(0.02), t(44)=3.69, p=0.009 | beta(SE)=0.15(0.03), t(44)=5.29, p=5.1e-05 | beta(SE)=0.16(0.04), t(44)=4.20, p=0.002 |
| Subcortical to Cortical SM | UCLA | beta(SE)=0.09(0.02), t(57)=3.81, p=0.005 | beta(SE)=0.11(0.04), t(57)=3.08, p=0.045 | beta(SE)=0.11(0.05), t(57)=2.53, p=ns |
| Subcortical to Cortical VI | CAMH | beta(SE)=0.06(0.02), t(103)=3.70, p=0.005 | beta(SE)=0.08(0.02), t(103)=4.05, p=0.001 | beta(SE)=0.08(0.03), t(103)=3.22, p=0.024 |
| Subcortical to Cortical VI | ZHH | beta(SE)=0.06(0.01), t(182)=4.46, p=2.0e-04 | beta(SE)=0.10(0.02), t(182)=5.08, p=1.3e-05 | beta(SE)=0.10(0.02), t(182)=4.81, p=4.5e-05 |
| Subcortical to Cortical VI | COBRE | beta(SE)=0.09(0.02), t(44)=3.79, p=0.006 | beta(SE)=0.12(0.03), t(44)=3.60, p=0.011 | beta(SE)=0.14(0.04), t(44)=3.75, p=0.007 |
| Subcortical to Cortical VI | UCLA | beta(SE)=0.03(0.02), t(57)=1.88, p=ns | beta(SE)=0.03(0.02), t(57)=1.75, p=ns | beta(SE)=0.05(0.02), t(57)=2.32, p=ns |
| Cortical DM to Cortical VA | CAMH | beta(SE)=-0.09(0.03), t(103)=-2.66, p=ns | beta(SE)=-0.07(0.02), t(103)=-3.23, p=0.023 | beta(SE)=-0.04(0.03), t(103)=-1.58, p=ns |
| Cortical DM to Cortical VA | ZHH | beta(SE)=-0.07(0.03), t(182)=-2.89, p=0.060 | beta(SE)=-0.08(0.02), t(182)=-5.13, p=1.0e-05 | beta(SE)=-0.09(0.02), t(182)=-3.67, p=0.004 |
| Cortical DM to Cortical VA | COBRE | beta(SE)=-0.16(0.05), t(44)=-3.25, p=0.031 | beta(SE)=-0.08(0.04), t(44)=-2.12, p=ns | beta(SE)=-0.13(0.05), t(44)=-2.76, p=ns |
| Cortical DM to Cortical VA | UCLA | beta(SE)=-0.12(0.05), t(57)=-2.40, p=ns | beta(SE)=-0.06(0.03), t(57)=-2.28, p=ns | beta(SE)=-0.10(0.04), t(57)=-2.65, p=ns |
| Cortical DM to Cortical DA | CAMH | NA | beta(SE)=-0.08(0.02), t(103)=-3.67, p=0.005 | beta(SE)=-0.06(0.03), t(103)=-2.25, p=ns |
| Cortical DM to Cortical DA | ZHH | NA | beta(SE)=-0.07(0.02), t(182)=-4.01, p=0.001 | beta(SE)=-0.09(0.02), t(182)=-4.61, p=1.1e-04 |
| Cortical DM to Cortical DA | COBRE | NA | beta(SE)=-0.07(0.04), t(44)=-1.61, p=ns | beta(SE)=-0.05(0.05), t(44)=-1.07, p=ns |
| Cortical DM to Cortical DA | UCLA | NA | beta(SE)=-0.07(0.02), t(57)=-2.92, p=0.071 | beta(SE)=-0.07(0.03), t(57)=-2.57, p=ns |
| Cortical FP to Cortical SM | CAMH | beta(SE)=0.06(0.02), t(103)=2.45, p=ns | beta(SE)=0.06(0.02), t(103)=3.10, p=0.035 | beta(SE)=0.08(0.03), t(103)=3.30, p=0.019 |
| Cortical FP to Cortical SM | ZHH | beta(SE)=0.10(0.03), t(182)=3.22, p=0.021 | beta(SE)=0.07(0.02), t(182)=3.94, p=0.002 | beta(SE)=0.08(0.02), t(182)=3.25, p=0.019 |

| | | | | |
|---|---|---|---|---|
| Cortical FP to Cortical SM | COBRE | beta(SE)=0.04(0.04), t(44)=1.18, p=ns | beta(SE)=0.10(0.04), t(44)=2.52, p=ns | beta(SE)=0.11(0.05), t(44)=2.17, p=ns |
| Cortical FP to Cortical SM | UCLA | beta(SE)=0.14(0.04), t(57)=3.54, p=0.011 | beta(SE)=0.06(0.03), t(57)=2.12, p=ns | beta(SE)=0.04(0.04), t(57)=1.10, p=ns |
| Cortical VA to Cortical DA | CAMH | beta(SE)=0.04(0.02), t(103)=1.88, p=ns | beta(SE)=0.05(0.02), t(103)=2.49, p=ns | beta(SE)=0.04(0.03), t(103)=1.27, p=ns |
| Cortical VA to Cortical DA | ZHH | beta(SE)=0.14(0.02), t(182)=5.58, p=1.2e-06 | beta(SE)=0.12(0.02), t(182)=6.86, p=1.5e-09 | beta(SE)=0.12(0.03), t(182)=4.69, p=7.5e-05 |
| Cortical VA to Cortical DA | COBRE | beta(SE)=0.06(0.03), t(44)=1.75, p=ns | beta(SE)=0.05(0.03), t(44)=1.67, p=ns | beta(SE)=0.06(0.04), t(44)=1.44, p=ns |
| Cortical VA to Cortical DA | UCLA | beta(SE)=0.09(0.04), t(57)=2.21, p=ns | beta(SE)=0.06(0.02), t(57)=2.71, p=ns | beta(SE)=0.11(0.02), t(57)=4.63, p=3.0e-04 |
| Cortical DA to Cortical SM | CAMH | beta(SE)=0.03(0.02), t(103)=1.65, p=ns | beta(SE)=0.07(0.02), t(103)=3.97, p=0.002 | beta(SE)=0.08(0.02), t(103)=3.46, p=0.011 |
| Cortical DA to Cortical SM | ZHH | beta(SE)=0.15(0.03), t(182)=4.87, p=3.3e-05 | beta(SE)=0.13(0.02), t(182)=6.08, p=9.9e-08 | beta(SE)=0.12(0.02), t(182)=4.95, p=2.4e-05 |
| Cortical DA to Cortical SM | COBRE | beta(SE)=0.05(0.04), t(44)=1.29, p=ns | beta(SE)=0.03(0.04), t(44)=0.75, p=ns | beta(SE)=0.06(0.04), t(44)=1.59, p=ns |
| Cortical DA to Cortical SM | UCLA | beta(SE)=0.07(0.04), t(57)=1.84, p=ns | beta(SE)=0.03(0.02), t(57)=1.08, p=ns | beta(SE)=0.02(0.03), t(57)=0.81, p=ns |
| Cortical VI to Cortical VI | CAMH | beta(SE)=-0.08(0.04), t(103)=-1.99, p=ns | beta(SE)=-0.12(0.04), t(103)=-3.09, p=0.036 | beta(SE)=-0.09(0.04), t(103)=-2.16, p=ns |
| Cortical VI to Cortical VI | ZHH | beta(SE)=-0.10(0.03), t(182)=-2.75, p=0.091 | beta(SE)=-0.15(0.03), t(182)=-4.65, p=8.8e-05 | beta(SE)=-0.15(0.03), t(182)=-4.26, p=4.7e-04 |
| Cortical VI to Cortical VI | COBRE | beta(SE)=-0.14(0.06), t(44)=-2.39, p=ns | beta(SE)=-0.11(0.05), t(44)=-2.08, p=ns | beta(SE)=-0.14(0.05), t(44)=-2.74, p=ns |
| Cortical VI to Cortical VI | UCLA | beta(SE)=-0.08(0.06), t(57)=-1.35, p=ns | beta(SE)=-0.05(0.04), t(57)=-1.11, p=ns | beta(SE)=-0.03(0.05), t(57)=-0.62, p=ns |
| Cerebellum to Striatum | CAMH | beta(SE)=-0.10(0.03), t(103)=-3.57, p=0.008 | beta(SE)=-0.10(0.03), t(103)=-3.57, p=0.008 | beta(SE)=-0.10(0.03), t(103)=-3.57, p=0.008 |
| Cerebellum to Striatum | ZHH | beta(SE)=-0.08(0.02), t(182)=-3.40, p=0.011 | beta(SE)=-0.08(0.02), t(182)=-3.40, p=0.011 | beta(SE)=-0.08(0.02), t(182)=-3.40, p=0.011 |
| Cerebellum to Striatum | COBRE | beta(SE)=-0.08(0.04), t(44)=-1.82, p=ns | beta(SE)=-0.08(0.04), t(44)=-1.82, p=ns | beta(SE)=-0.08(0.04), t(44)=-1.82, p=ns |
| Cerebellum to Striatum | UCLA | beta(SE)=-0.09(0.03), t(57)=-2.96, p=0.062 | beta(SE)=-0.09(0.03), t(57)=-2.96, p=0.062 | beta(SE)=-0.09(0.03), t(57)=-2.96, p=0.062 |

| | | | | |
|---|---|---|---|---|
| Cerebellum to Thalamus | CAMH | beta(SE)=-0.13(0.03), t(103)=-4.62, p=1.6e-04 | beta(SE)=-0.13(0.03), t(103)=-4.62, p=1.6e-04 | beta(SE)=-0.13(0.03), t(103)=-4.62, p=1.6e-04 |
| Cerebellum to Thalamus | ZHH | beta(SE)=-0.10(0.02), t(182)=-4.61, p=1.0e-04 | beta(SE)=-0.10(0.02), t(182)=-4.61, p=1.0e-04 | beta(SE)=-0.10(0.02), t(182)=-4.61, p=1.0e-04 |
| Cerebellum to Thalamus | COBRE | beta(SE)=-0.11(0.04), t(44)=-2.67, p=ns | beta(SE)=-0.11(0.04), t(44)=-2.67, p=ns | beta(SE)=-0.11(0.04), t(44)=-2.67, p=ns |
| Cerebellum to Thalamus | UCLA | beta(SE)=-0.15(0.03), t(57)=-4.73, p=2.1e-04 | beta(SE)=-0.15(0.03), t(57)=-4.73, p=2.1e-04 | beta(SE)=-0.15(0.03), t(57)=-4.73, p=2.1e-04 |

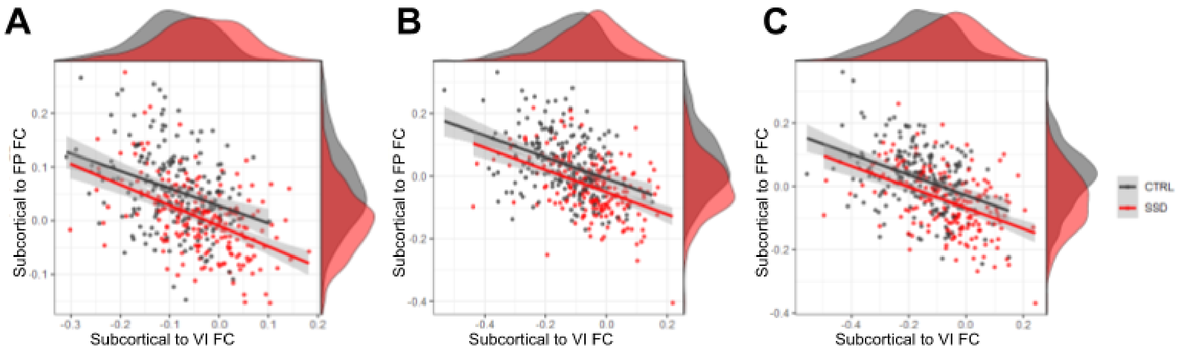


Supplemental Figure 2. Correlation between subcortical to VI functional connectivity with subcortical to FP functional connectivity in participants with SSD (red) and Healthy Controls (HC). The correlation is present for the( A) volume-based (B), surface-based and (C) personalized surface-based (PINT) approaches.
